# Metagenomic insights into ecosystem function in the microbial mats of Blue Holes, Shark Bay

**DOI:** 10.1101/2020.09.18.304444

**Authors:** Gareth S Kindler, Hon Lun Wong, Anthony W D Larkum, Michael Johnson, Fraser I MacLeod, Brendan P Burns

## Abstract

Microbial mat ecosystems vary in complexity and structure depending on the environmental constraints placed by nature. Here, we describe in detail for the first time the community composition and functional potential of the microbial mats found in the supratidal, gypsum-rich, and hypersaline region of Blue Holes, Shark Bay. This was achieved via high throughput sequencing of total mat community DNA on the Illumina NextSeq platform. Mat communities were mainly comprised of Proteobacteria (29%), followed by Bacteroidetes/Chlorobi Group (11%), and Planctomycetes (10%). These mats were found to also harbor a diverse community of potentially novel microorganisms including members from the DPANN and Asgard archaea, Candidate Phyla Radiation (CPR) and other candidate phyla, with highest diversity indices found in the lower regions of the mat. Major metabolic cycles belonging to sulfur, carbon, nitrogen, and fermentation were detected in the mat metagenomes with the assimilatory sulfate reduction pathway being distinctly abundant. Critical microbial interactions were also inferred, and from 117 medium-to-high quality metagenome-assembled genomes (MAGs), viral defense mechanisms (CRISPR, BREX, and DISARM), elemental transport, osmoprotection, heavy metal and UV resistance were also detected in the mats. These analyses have provided a greater understanding of these distinct mat systems in Shark Bay, including key insights into adaptive responses.

## Introduction

Microbial mats are vertically stratified organosedimentary aggregations, embedded in a matrix of minerals, growing in often extreme habitats [1, 2]. Vertical stratification of microbial mats is attributed to physicochemical gradients, which favour niche differentiation between microorganisms [3]. The microbial community is a diverse mixture of bacteria, archaea, eukarya and viruses [4, 5], which vary taxonomically and functionally depending on the environment in which the mat forms. These microorganisms perform a range of functional roles, with nutrient cycling capabilities spanning across major elements of carbon, nitrogen, and sulfur [6], resulting in a self-sustaining community, highlighted by numerous microbial interactions.

Modern microbial mats are often considered analogues of ancient stromatolites, the fossilised communities of 3.5 billion years past [7]. During this time Earth was experiencing significant biogeochemical transitions and extremes [8]. Precambrian stromatolites, lithified laminated formations represent the oldest ecosystems known, and have played a crucial role in oxygenation of earth’s atmosphere, enabling higher forms of life to evolve [9, 10]. The same lithification mechanisms are extremely rare to occur in present day microbial mats, but Late Archaean Earth harboured similar microbial mat communities [11]. Microbial mats and stromatolites are considered as major influencers on global biogeochemistry and evolutionary processes for half of Earth’s history. Modern microbial mats continue to contribute to increased biological productivity, shared reducing power, and oxidation of Earth’s surface [12]. Distinct phyla of archaea found in modern microbial mats have recently been suggested to be the origin of eukaryotic cellular complexity [6, 13].

Microbial mats are exposed to conditions thought to restrict life such as temperature, solar radiation, desiccation, and salinity. Studies have shown modern microbial mats to be prevalent in unique locations across the globe such as karstic spring mounds [14], hot springs and geysers [15], hypersaline and volcanic lakes [16-19], rivers [20], acid mine drainages [21] and supratidal pools [22]. In the intertidal and subtidal hypersaline marine environments of Shark Bay, Western Australia [23, 24], modern microbial mats and stromatolites continue to thrive as one of the best examples in the world.

On the coast of Western Australia, two arms of the sea come together to form the barred basin of Hamelin Pool (Figure S1). While the microbial mats of Hamelin Pool have been the subject of recent intense research [6, 24, 25], the distinct ecosystem of Blue Holes is uncharacterized. Located on the Nanga Peninsula, on an embayment of Hamelin Pool there are series of hypersaline ponds known as Blue Holes (Figure S2). The site consists of 12 circular holes on a 650 m wide supratidal flat, with each hole having a depth of up to 2.5 m and diameter between 2 to 30 m [25]. Between 900 years ago and present day sea levels decreased by 1.5 m [26], during this time it is likely the tidal flat opened up, combined with the groundwater dissolution of fossilised gypsum (CaH4O6S) it resulted in the formation of Blue Holes [25]. Blue Holes is located in the now supratidal zone with periodic recharges of rainwater, abnormal high tides from Hamelin Pool and prolonged on-shore winds [26, 27] creating a surface water layer of lower salinity, density, and temperature than the underlying hypersaline water layer [25]. Stratification of the water column is an intermittent feature of these holes [27], as periods of low water levels, wind action, and inundation during very high tides can diminish stratification (Figure S3). An early study in 1980 identified microbial mats of salmon pink and blue-green pigments covering the sedimentary base of Blue Holes [27]. A high level of microorganism diversity was observed with two dominant cyanobacteria thought to be responsible for the mat architecture, Phormidium sp. and Aphanothece sp. Recently in 2014, water from Blue Holes was measured to have a salinity of 78 Practical Salinity Units (PSU), double that of normal seawater, and to be sulfidic [5]. Both interactions were preliminary and mainly served to add to the intrigue surrounding the holes.

Although important characteristics of the Blue Holes ecosystem have been described, the abiotic and biotic nature at the molecular scale remains to be illuminated. Previous work on microbial mats from another site in Shark Bay (Nilemah, Figure S1) has found extensive diversity, contributing to understanding the roles of biogeochemical cycling, adaptations, and novel pathways and microorganisms in these ecosystems [6, 28]. For the first time, we present detailed analyses of microbial community structure and function in the mats of Blue Holes, Shark Bay using high throughput metagenomic sequencing.

## Methods

### Location Description and Sampling, DNA Isolation, Sample Sectioning

Microbial mat and water samples were collected from Blue Holes (Hamelin Pool, Shark Bay, WA, 26°15’39.6”S 113°56’57.3”E) (Figure S2) on the 30th of October 2017. Of the three most discrete holes, the smallest was sampled. Using a sterile metal spatula, microbial mat slabs of 10 cm^2^ were cut out 1 m from the edge of a pool and at an approximate water depth of 20 cm (Figure S2). Samples were placed in RNAlater solution. Water overlying the Blue Holes microbial mats was collected along with water from Nilemah (Hamelin Pool, Shark Bay, WA, 26°27’336’’S, 114°05’762’’E), for reference (Figure S1). Separation of the vertically stratified Blue Holes microbial mat was based on visible pigmentation along a depth gradient (Figure S2). The sample was sliced longitudinally to form three equally sized layers (~ 7 mm in depth). The surface layer (green) comprised the first 7 mm of microbial mat, subsurface (pale red) was designated 7 to 14 mm, and base (translucent) as 14 to 21 mm (Figure S2).

### Nucleic Acid Extraction and Sequencing

Extraction of nucleic acids from microbial mat samples was performed with a PowerBiofilm DNA Isolation Kit (10). Weight of starting microbial mat material was between the 0.16 and 0.20 g for all extraction samples. Nucleic acids were extracted in triplicate from each layer of microbial mat, resulting in a total of nine samples (Figure S4). Extractions were undertaken following the manufacturer’s instructions, with several modifications. During the mechanical bead beating step, the PowerBiofilm Bead Tube was homogenised using a Fastprep24 instrument at a speed of 5.5 m/s for a 30 s duration. The volume of BF3/IRS Solution was increased from 100 μL to 200 μL to aid in the removal of inhibitors such as exopolymeric substances (EPS). All 60 s centrifugation steps were increased to 90 s, excluding the step involving the selective binding of DNA to the silica membrane. An additional 60 s of centrifugation was also added to the removal of residual wash and release of DNA from Spin Filter membrane steps. To further purify the extracted nucleic acids, an ethanol precipitation procedure was employed. To each tube containing suspended DNA, 50 μL of 7.5 M Sodium Acetate and 500 μL of −20°C 100 % ethanol was added. Tubes were incubated at −20°C for 24 h. Following incubation, tubes were centrifuged for 30 min at 14,600 × g. The supernatant was then removed by pipetting. To wash and resuspend the DNA, 500 μL of 80 % ethanol was added. Tubes were centrifuged at 14,600 × g for 15 min. The wash step was repeated, followed by the supernatant being pipetted out, ethanol added to resuspend the DNA, followed by centrifugation for 15 min. Supernatant was pipetted out and tubes were left to air dry. DNA was resuspended in 100 μL of BF7/EB. Tubes were incubated on wet ice for 30 min, then stored at −20°C. A Nanodrop ND-1000 Spectrophotometer was used to determine DNA quality and quantity according to the manufacturers protocol. To check for degradation sample quality was visualised through agarose gel electrophoresis. Ramaciotti Centre for Genomics (UNSW Sydney, Australia) performed the library preparation and metagenomic sequencing. Libraries were prepared using the Nextera XT DNA Sample Preparation Kit, with an input of 1 ng of DNA per sample. A final clean-up step at a ratio of 0.7 bead volume to 1 pooled library volume was performed to eliminate larger fragments of an abnormal sample library. Illumina Nextseq 500 with a run format of Medium Output 2×150 bp was used to sequence the libraries. Prior to assembly, all biological replicate sequence files (three from each microbial mat layer) were concatenated to create pooled files from each layer of microbial mat. Quality checking of nucleic acid reads was performed by FastQC (70). Trimmomatic (71) was used to cut away low quality bases using a sliding window setting of 4:21 and minimum length of 50 bps (Figure S4).

### Metagenome analyses

Contig assembly for metagenome analyses was performed using the meta option through SPAdes [29]. Metagenome statistics were generated using QUAST [30]. Prediction of amino acid sequences was completed using Prodigal [31]. For taxonomic classification, 40 marker genes or COGs corresponding to single-copy gene families universally distributed in prokaryotic genomes [32] were extracted using FetchMGs [33] (https://github.com/motu-tool/fetchMGs). We compared our single-copy gene sequences with the non-redundant protein database (ftp://ftp.ncbi.nlm.nih.gov/blast/db/FASTA/ on the 22nd May) - RefSeq Release 94 (May 17, 2019) [34], through the DIAMOND software (diamond v0.9.24.125) [35] (Figure S4). Only the best hit of each gene was retained, using a minimum amino acid identity of 50% over at least 80% of the query length [36]. Using an adhoc script in PyCharm, through utilizing the Pandas [37] and ETE 3 [38] packages, sequence Tax IDs were converted into counts for each marker gene. Marker gene counts were converted through a compositional normalization process to relative abundances. Using ggplot2 (https://ggplot2.tidyverse.org/) in R software (http://cran.r-project.org/), the relative abundances were visualized on a stacked bar chart.

Metagenomes were annotated for both function and taxonomy using the KEGG online server [39]. Hits were kept using a GHOSTX cut-off of 100. Using MUSiCC (Metagenomic Universal Single-Copy Correction), gene abundances were normalized [40]. In accordance with previous work, genes unique to a pathway or cycle were considered to be diagnostic of that metabolism and were averaged to calculate the metabolic abundances [6, 36, 41, 42]. Using ggplot2, stacked bar charts containing fill correlated the KEGG-annotated taxonomy with diagnostic genes.

### Metagenome-assembled genomes

Contig assembly was performed using Megahit [43] (Figure S4). The minimum k-mer length was set to 27 and incremental k-mer size to 10. Resulting contigs were then subsampled for lengths greater than 1000 bps, conditions which are a required for downstream functional annotation. Indexing of reads was performed using BWA [44]. BWA mem assembled sequences were mapped against the previously trimmed sequences. View and sort functions in SAM Tools was then used to compact files into the practical bam format [45]. Bam files are used in combination with assembled contig files for downstream analyses.

Input coverage and depth files for binning programs were produced using standard program settings. To generate genomes, contigs were grouped into bins using MetaBAT [46], Maxbin2 [47] and Concoct [48]. Using the three sets of binning results, optimised and non-redundant bins were calculated using DAS Tool [49], resulting in bins of improved quality. Refined bins, now referred to as metagenome-assembled genomes (MAGs) were used for all downstream analysis. Estimation of quality measurements of MAGs was performed using CheckM [50], on default settings. Quality of genomes were then categorised using the standards of Minimum Information about Metagenome-Assembled Genomes (MIMAG) [51]. MAGs with greater than 10% contamination were discarded. To survey MAGs for the presence of subunit rRNA genes, RNAmmer [52] was used. However, due to incongruent results the rRNA criterion of the MIMAG standards was exempted from this study, conforming with previous work [6]. Identification of the amount and types of tRNA sequences was performed using tRNAscan-SE.

Due to their ubiquity across bacterial and archaeal genomes, 15 specific ribosomal proteins were chosen for phylogenomic based analyses (ribosomal proteins L2, L3, L4, L5, L6, L14, L15, L18, L22, L24, S3, S8, S10, S17, and S19) [53]. MAGs containing greater than six ribosomal protein sequences generated by Phylosift and of low, medium, and high quality were used for construction of the phylogenetic tree. If ribosomal sequence files contained more than one sequence, the shorter was deleted. Sequences were aligned using MAFFT [54], followed by BMGE [55] to remove gaps within sequences. The phylogenetic tree was constructed by maximum likelihood using IQ-TREE [56]. A total of 1000 bootstrap replicates was used to infer confidence in the final phylogenetic tree construction. Branches with bootstrap values lower than 50 were deleted in the Interactive Tree of Life online interface (iTOL) [57]. Using CheckM, Phylosift [58] and NCBI Protein BLAST [59] MAGs were assigned taxonomy and annotated onto the formatted phylogenetic tree in Inkscape [60] (Table S1). Within NCBI Protein BLAST, a threshold of greater than 40% aminoacid identity and cut-off E-value of 1e-20 was used.

Contigs of nucleotide sequences were converted to protein-coding open reading frames (ORFs) using Prodigal, within CheckM (Table S1). To annotate protein functions of MAGs, a combination of KEGG server [39] and InterProScan [61] was used. For KEGG annotation, genes were recorded as present using a KEGG confidence score of greater than 85 [6]. Genes annotated by InterProScan were considered present using a cut-off value of lower than 1e-10. Annotated protein sequences of MAGs were compared across a range of genes involved in metabolisms and adaptive responses. Metagenome-assembled genomes of >70% completeness [62] and <10% contamination (medium and high quality) were used for metabolic potential analyses and corresponding phylogenomic analyses. Statistical analysis of MAGs was performed using the R software. Figure 3 was built in R using the ggplot2 package. Genes related to foreign genetic element defence were recorded as either present, partially present, or absent. Genes related to environmental adaptions were recorded as either present or absent. Phylogenomic tree attached to Figure 3 was created despite two MAGs containing less than six ribosomal proteins.

### Water chemistry

To determine cation, dissolved nitrogen, and dissolved organic carbon concentrations of water samples the following analyses were conducted in triplicate at the Mark Wainwright Centre (UNSW Sydney, Australia). Inductively coupled plasma mass spectrometry (ICP-MS) and liquid chromatography with organic carbon detection (LC-OCD) were used for these analyses. Salinity measurements were undertaken with a refractometer (Bellingham & Stanley). Water samples were filtered through a 0.45 μm syringe driven unit (Millipore). Water was sampled from two separate microbial mat sites in Shark Bay (Blue Holes and Nilemah) and measured in duplicate. Redox potential and pH measurements were undertaken using a pH Cube (TPS).

## Results

We studied the ecosystem of Blue Holes, Shark Bay, focusing our analyses on the microbial mat and overlying water. Rainfall in Shark Bay over the month preceding collection (October 2017) was dry, only receiving 1 mm. The average of the month’s daily solar radiation was 25.55 MJ m-2 with daily temperatures ranging between 15 and 28°C (Table S2). Blue Holes water salinity was 109.8 PSU, sodium (31634 mg/L) was the most abundant cation, followed by magnesium (3804 mg/L) (Table S3). Sulfur (2983 mg/L) is the third most abundant cation in the Blue Holes ecosystem, 151% higher than Nilemah. Dissolved Organic Nitrogen (DON) and Dissolved Organic Carbon (DOC) were measured at 19.7 mg/L and 11.1 mg/L, respectively. The Blue Holes microbial mat samples were sectioned along three distinct layers: a surface layer of green and translucent and filamentous properties (0-7 mm), a subsurface layer of maroon and green (7-14 mm), and a base layer saturated by sediment and Fragum erugatum shells (translucent, 14-21 mm) (Figure S2). We generated metagenomes from concatenating triplicate sequenced samples for each layer of microbial mat. Each metagenome comprised a GC content greater than 55% (Table S4).

### Community composition

To characterize the microbial composition of the Blue Holes mats, we used the occurrence of universally distributed, single-copy marker genes within each metagenome or layer (Table S5). Based on the assignment of taxonomy to these genes, we detected a range of microbial diversity within the layers of microbial mat as shown through Shannon entropy (2.3-3.1) and Gini-Simpson (0.8-0.9) indices (Table S4). The surface layer was the least diverse of the three, and as depth increased so did diversity resulting in the base layer being twice as diverse as the surface. The middle layer, or subsurface showed a level of diversity similar to the base with 12% less. At the surface layer, bacteria dominated with representation of Proteobacteria, the Bacteriodetes/Chlorobi group, Planctomycetes, and Spirochaetes (Figure 1). Although in different orders of prevalence, these four phyla are represented as the most dominant throughout each mat layer. Cyanobacteria were present within the mat, albeit at low levels (Table S6). Proteobacteria is the most dominant phylum in each layer of the mat, and accounts for the most in the surface at 38% of all single-copy genes in the metagenome. The majority of bacterial candidate phyla were present in abundances less than 1% of the total metagenome sequences. However, cumulatively they accounted for 7%, 13%, and 15% from the surface to base microbial mat layers, respectively. Of the bacterial candidate phyla, 49 phyla accounted for 95-97% of the diversity across the three microbial mat layers (Figure S5a/Table S6) whilst a total of 105 were detected. Candidatus Omnitrophica dominated the upper two layers, whilst Candidatus Zixibacteria dominated the lower two layers. Candidatus KSB1 and Candidatus Falkowbacteria followed in abundance in the base layer. As depth increases, the prevalence of bacteria decreases, from 98% in the surface to 92% in the base, with archaea filling the space (Figure S5b/Table S6). Woesarchaeota and Euryarchaeota were profuse throughout the mat layers, following opposing trends along the depth gradient. In smaller quantities, Aenigmarchaeota, Altiarchaeota, and Bathyarchaeota were present. Of the Asgard archaea, Lokiarchaeota outshone Odinarchaeota and Thorarchaeota to be the most prevalent.

**Figure 1.**
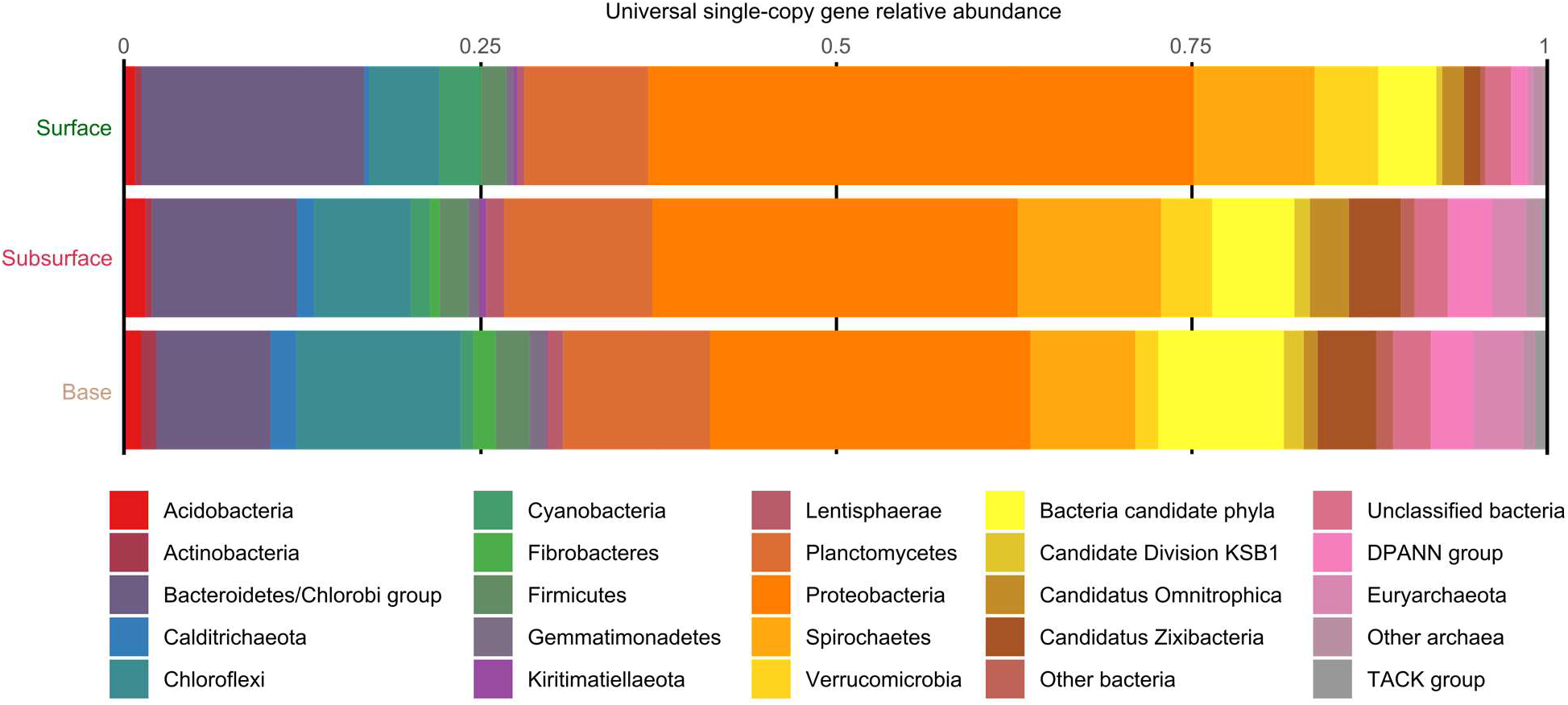
Overall microbial relative abundance within the microbial mat metagenomes of Blue Holes (Shark Bay, Australia), as inferred from the taxonomic assignment of 40 universally distributed single-copy genes (COGs). Candidate Division KSB1, Candidatus Omnitrophica, and Candidatus Zixibacteria are separate of their respective phyla or other clades. Each metagenome is the result of merging the three biological replicates, respectively. Further classification of bacterial and candidate phyla is shown in Figure S5.

### Metabolic cycles and pathways

We correlated MUSiCC-normalised gene abundances of targeted metabolic cycles and pathways with taxonomy to elucidate the likely metabolic roles different phyla are playing within the mat layers of these ecosystems (Figure 2). Genes unique to a specific metabolic process were considered diagnostic (Table S7) in accordance with previous research [36]. Sulfur cycling was the most abundant metabolism investigated, followed by carbon and nitrogen (Table S8), while methanogenesis accounted for the least number of genes. Of all metabolic pathways, assimilatory sulfate reduction genes were found to be the most prevalent. Fermentation was the next abundant, followed by dissimilatory sulfate reduction and oxidation. The reductive acetyl-CoA pathway dominates carbon metabolisms, followed by the reductive pentose phosphate cycle, DC-HB/HP-HB cycles, 3-HP bicycle, and the reverse citric acid cycle (rTCA). Within the nitrogen cycling capabilities of the mat, nitrogen fixation was the most prevalent, followed by dissimilatory nitrate reduction. Prominent nitrogen fixers within the mat were Proteobacteria, Chloroflexi, Cyanobacteria, and Verrucomicrobia, shifting in abundance depending on the layer. The least number of genes within the mat were associated with nitrification. Proteobacteria account for 10% of all metabolic genes investigated in this study. Deltaproteobacteria (24%) are responsible for the largest average portion of combined metagenome genes across all metabolisms, followed by Alphaproteobacteria (12%). Gammaprotoebacteria, Bacteroidetes, Firmicutes, Planctomyctes account for 33% of metabolic genes (Table S8). Eukarya genes were identified to be present within carbon, nitrogen, phototrophy, sulfur, fermentation, and oxidative phosphorylation metabolisms. Anoxygenic photosynthesis genes, mostly contributed by those associated with bacteriochlorophyll biosynthesis, were five times more abundant than oxygenic photosynthesis genes. Deltaproteobacteria, Chloroflexi, and Firmicutes contributed to anoxygenic photosynthesis (BC), whereas oxygenic photosynthesis was associated with Planctomycetes and a diverse selection of other phyla. Prominent cascades are found in nitrogen fixation and anoxygenic photosynthesis, showing the highest gene occurrences to be within the surface mat layer (Figure 2). Inverse cascades can be seen in DSRO and the reductive acetyl-CoA pathway. Most nitrogen-associated pathways were relatively evenly represented across the mat layers. The mat layer with most diagnostic metabolic genes was the base, followed by the surface and subsurface. Methanogenesis genes are the lowest in surface and increase slightly with depth.

**Figure 2.**
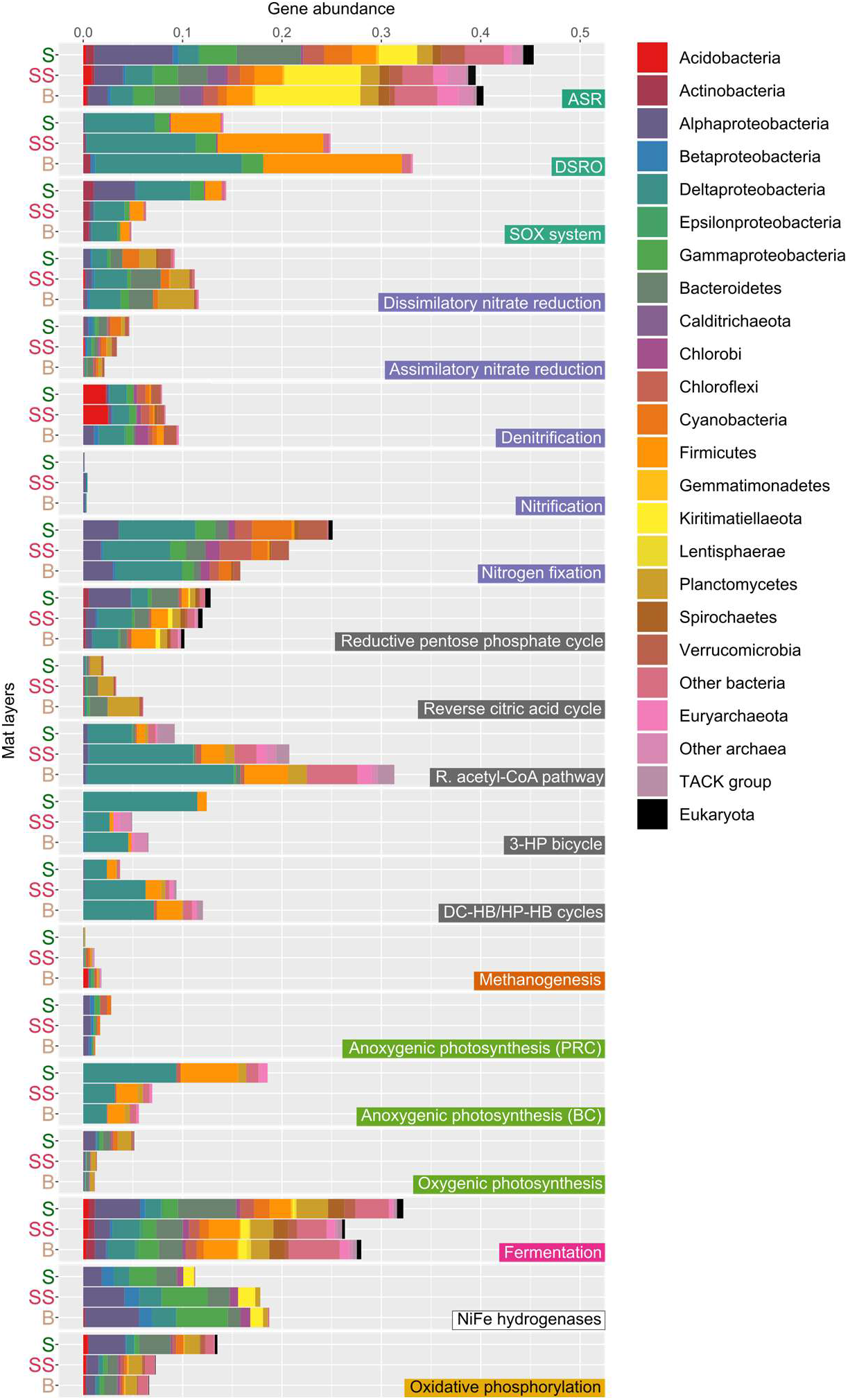
Metabolism and associated taxonomy in Blue Holes microbial mats. Stacked bar histograms showing taxonomic classification of the KEGG-annotated metabolic MUSiCC-normalised gene abundances of the three layers of microbial mat (S, surface; SS, subsurface; B, base). ASR, Assimilatory sulfate reduction; DSRO, Dissimilatory sulfate reduction and oxidation; Reductive pentose phosphate cycle (Calvin-Benson-Bassham cycle); Reverse citric acid cycle (Arnon-Buchanan cycle); R. acetyl-CoA pathway, reductive acetyl-CoA pathway (Wood-Ljungdahl pathway); Dicarboxylate-hydroxybutyrate (DC-HB) and Hydroxypropionate-hydroxybutylate (HP-HB) cycles; Anoxygenic photosynthesis (PRC), photosystem reaction center in purple bacteria; Anoxygenic photosynthesis (BC), Bacteriochlorophyll biosynthesis in green non-sulfur.

### Environmental Adaptations

A total of 156 metagenome-assembled genomes (MAGs) of low, medium, and high-quality were uncovered from the binning process. MAGs of greater than 70% completeness and less than 10% contamination were included in comparative genomic analysis, constituting a total of 117 (Figure 3). Defence systems protecting against foreign genetic elements are present within the Blue Holes microbial mat MAGs. CRISPR-associated (Clustered Regularly Interspaced Short Palindromic Repeats) or cas proteins were identified in a variety of bacterial and a single Euryarchaeota (BH_18) MAG(s) (Figure 3). The microbial defence system, Defence Island System Associated with Restriction–Modification (DISARM) (105) genes were identified to partially present in 48% of bacterial MAGs (complete presence in 39%). Forty-one MAGs contained protein domain DUF1998 (drmB PF09369) located on the same contig as protein domain drmA (PF00271), indicating community members may harbour the DISARM system (105) (Table S9). Core genes (pglZ, brxC/pglY) of the recently discovered, Bacteriophage Exclusion (BREX) system were detected in partial presence across 38% of bacterial MAG phyla (106).

**Figure 3.**
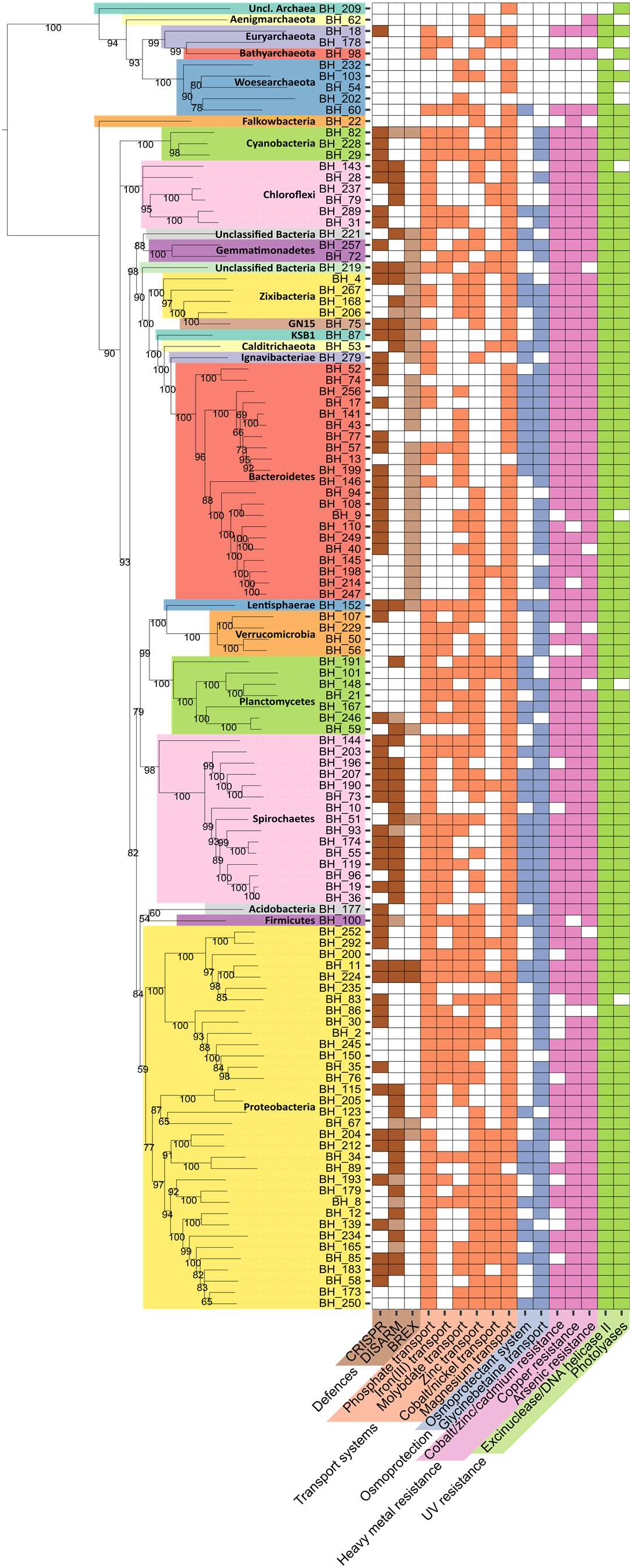
Colour differentiated phylum-level phylogenomic tree and table displaying MAG functionality. X-axis indicates pathways and genes implicated in environmental adaptions. The longitudinal lines of the table correspond to MAG labels located on the y-axis. Connected to the y-axis is a phylogenetic tree created from MAG ribosomal proteins, showing bootstrap values. White squares indicate the absence of a given pathway in a MAG. Abbreviations: clustered regularly interspaced palindromic repeats (CRISPR), defence island system associated with restriction–modification (DISARM), and bacteriophage exclusion (BREX) system.

Of transport system genes, phosphate and magnesium transporting genes were the most abundant in MAGs. Genes related to osmoprotection (opuABDC) and glycine betaine transport (opuD, bet, proVWX) were detected in 85% of MAGs (Table S9). The presence of heavy metal cations in Blue Holes water is complemented by the high representation of heavy metal resistance genes across MAGs. Cobalt-zinc-cadmium (zntA, czcABCD), copper (copAB, cutC, cusAB), arsenic (ARSC12, arsAH, arsB/acr3) genes were present in most bacterial MAGs but are mostly lacking within the archaea (Figure 3). Despite this, a Bathyarchaeota (BH_98), Woesearchaeota (BH_60) and Euryarchaeota (BH_18) MAG contain every gene related to all three heavy metals investigated. Genes associated with resistance to harmful UV light were ubiquitous across all MAGs (Figure 3). MAGs contained genes encoding for either an exinuclease (uvrABC), DNA helicase II (uvrD), spore photoproduct lyase lyase (splB), or deoxyribodipyrimidine photo-lyase (phrB).

## Discussion

The microbial mat systems that dominate Shark Bay have been extensively studied, however primarily at the site of Nilemah in Hamelin Pool notably Nilemah’s [3, 6, 63]. Prior to the present study the intriguing ecosystem of Blue Holes has remained unexplored. We present, for the first time, an investigation of Blue Holes at the genetic and chemical level. Blue Holes is an ecosystem governed by constant fluctuation. It experiences an overall salinity range of at least 78-109.8 PSU [5], in context, the neighboring microbial mat site of Nilemah is consistently exposed to a salinity range of 55-70 PSU [25], double the salinity of open ocean water (Table S3). When taken together these results indicate the nature and location of the holes make them more vulnerable to the diel, seasonal, and intermittent weather changes than the broader Shark Bay ecosystem (Figure S3). As harsh conditions arise (prolonged dry periods, storms, or powerful tides), Blue Holes microorganisms are likely either being pushed into dormancy [64] or dying off. This is followed by a period of stability in which the microorganisms are resuscitated or re-seeded from the transfer of sediment or water from surrounding holes or the larger Hamelin Pool (Figure S1).

The Blue Holes microbial mat harbors a diverse community of microorganisms (Figure 1). Proteobacteria, the Bacteriodetes/Chlorobi group, Planctomycetes and Spirochaetes are the dominant phyla across each mat layer. Although the prevalence of Proteobacteria and the Bacteroidetes/Chlorobi group are shared features with the Nilemah mats of Shark Bay [65], abundances of the other phyla greatly differ. The most diversity in the Blue Holes mats is in the lower mat layers, further away from the overlying water and atmosphere (Table S4). The increasing diversity indices along the depth gradient is driven by the greater share of bacterial candidate phyla and archaea in the lower layers (Table S6). While most bacterial candidate genes were accounted for by 49 phyla, more than double that amount was detected. Of those identified in this study a significant portion belong to the Candidate Phyla Radiation (CPR), such as Candidatus Falkowbacteria. Along with CPRs, the DPANN superphylum of archaea detected in this study contribute to the microbial dark matter (MDM) in Blue Holes mats. Three subgroups belonging to the Asgard archaea were detected in this study and when taken together with findings from the Nilemah mats [6] are beginning to illuminate the share of diversity MDM has in the Shark Bay systems. Although higher average GC content of communities has been shown to be potentially indicative of increased complexity and competition [66], trends between environments are highly variable [67]. The GC values obtained within this study were higher than average (>55%) (Table S4), similar to other microbial mats [36].

Delta-, Alpha-, Gammaprotoebacteria, Bacteroidetes, Firmicutes, Planctomyctes most likely account for most of the primary production in Blue Holes mats. Individually, deltaproteobacteria are the driving force across the investigated metabolisms, accounting for the highest number of detected genes (Table S8). The remaining microorganisms are likely heterotrophic and obtain their energy through the remaining material of organic or inorganic cascades [36]. Recent observations of the Blue Holes environment noted the water to smell strongly sulfidic [5], and contain the sulfate evaporite, gypsum [25]. Sulfur is the third most abundant cation in the Blue Holes water (Table S3) and within the function of the mat, sulfur cycling, specifically assimilatory sulfate reduction is the most prevalent metabolism showing little disparity between layers as it is anoxygenic (Figure 2). Blue Holes as an ecosystem seems to mimic the sulfate evaporite basins that appeared with the Great Oxidation Event (GOE) [68]. Like early environments, autotrophs in oligotrophic environments such as Blue Holes may be at an advantage by assimilating abundant molecules such as atmospheric CO_2_ into cell material. Of the six carbon-based metabolisms, the strictly anaerobic reductive acetyl-CoA pathway is the most prevalent within our mats (Figure 2). Deltaproteobacteria, Firmicutes, the TACK group are the key drivers of this C fixation pathway. Claims suggestive of the acetyl-CoA pathway being one of the oldest carbon fixing pathways [69] and along with the rTCA being ancestral [70], support the premise of modern microbial mats as windows to the past [36, 63]. The reductive acetyl-CoA pathway, but also the dicarboxylate-hydroxybutyrate (DC-HB), hydroxypropionate-hydroxybutylate (HP-HB) cycles, and the rTCA cycle showed inverse cascades along the depth gradient. Methanogenesis genes were in relatively minute amounts but increased with depth, indicating methanogenesis could be occurring deeper in the mats than examined here [8]. The mat layers most exposed to oxic conditions harboured significantly higher gene counts of SOX system, nitrogen fixation, 3-HP, and oxidative phosphorylation (Figure 2). Along with oxygenic photosynthesis, anoxygenic photosynthesis genes were more abundant in the oxic layers than the anoxic zone, a similarity shared with microbial mat ponds in the Atacama Desert [36].

Biotic stressors within the Blue Holes ecosystem can come in the form of foreign genetic elements, revealed to play a major role in Shark Bay microbial mats (15), potentially impacting ecosystem function. As part of the CRISPR-Cas immune system, CRISPRs and CRISPR-associated (Cas) proteins function to provide an RNA-guided defence mechanism against viruses, plasmids, and transposable elements [71]. Within Blue Holes, protection against harm from foreign genetic elements is present in 55% of bacterial MAGs, and 10% of archaeal MAGs (Figure 3). This uneven distribution is inverse to current observations which have identified CRISPR-Cas systems in approximately 50% of bacterial genomes and 90% of archaeal genomes [72]. The two novel defence systems DISARM [73] and BREX are widespread in Blue Holes bacterial MAGs and no presence in the archaeal MAGs. The inverse skew could be explained by the relative paucity of archaeal genomes or suggested complementarity and heterotrophic lifestyles of phyla such as Woesearchaeota [74].

Abiotic factors such as solar radiation, high elemental concentrations, fluctuating salinity, and temperature ranges (Table S2/S3)) contribute to the stressors in the Blue Holes ecosystem. In the overlying water, magnesium is the most abundant cation after sodium, and it follows that genes encoding for a magnesium transporter (mgtE) are among the highest represented transport genes in MAGs (Table S9). A total of 47% of Blue Holes MAGs encode for both arsenate reductase (arsC) and an arsenate transporter (acr3), indicating potential protection against arsenic [76], albeit they are in low levels in Blue Holes water. Through the identification of ‘arsenic-rich’ regions of fossilised stromatolites, arsenic metabolism has been proposed to have a role in early earth environments belonging to more than 3.4 million years ago [75]. This suggests genes responsible for arsenic metabolism in Blue Holes could be ‘carried on’ from periods such as the Archaean, when arsenic was in relatively high abundance. Cations of copper, cobalt, zinc, and cadmium share this similarity of being minimal in Blue Holes water (Table S3) while occurrence of genes related to movement of these cations across membranes in MAGs are prevalent. The accumulation of osmolytes as an adaptive mechanism to high osmolarity environments is evolutionarily conserved within bacteria and archaea, and highly represented in Blue Holes MAGs (77%). The occurrence of genes encoding for transporters of glycine betaine and proline (opuD, bet, proVWX) is greater than the Nilemah, Shark Bay microbial mats (28% of MAGs), which are subjected to less salinity stress. To combat UV harm the microbial mats of Blue Holes were found to be equipped with genes encoding for excinucleases (uvrABC), DNA helicase II (uvrD), spore photoproduct lyase (splB), and deoxyribodipyrimidine photo-lyase (phrB) (Table S9). Excinuclease proteins [77] and helicase II enzymes [78] function to cleave damaged DNA, and are encoded in 98% of MAGs, making them the most abundant set of genes in all MAGs.

### Conclusions

This study described for the first time at millimetre resolution the microbial community structure and function of the mat ecosystems of Blue Holes, Shark Bay using a metagenomic approach. While metagenomics provides an excellent platform for understanding microbial systems, future analyses at the level of gene or protein expression (such as metatranscriptomics and metaproteomics), or even single cell genomics will provide additional scaffolding to confirm the interactions and major pathways inferred here.

## Supporting information

https://drive.google.com/drive/folders/1hdAbW-tnwMIu5FRgziQOsf89n0OJ2sHQ?usp=sharing

## Acknowledgments

We thank the ResTech team from UNSW for sharing their time and knowledge regarding everything computers. Additionally, Josh Hamm for his welcome advice and discussion surrounding the bioinformatic process and Tim Williams for his willingness to share his knowledge. We thank Sarah Westcott from mothur for her help and Alessio Milanese from EMBL for his help with metagenomics.

## Author contributions

G. S. Kindler performed sample preparation, data handling, data analyses, and wrote the manuscript. H. L. Wong supervised and assisted with wet and dry laboratory work, along with proof-reading. A. Larkum, M. Johnson and F. Macleod contributed to specific analyses. B. P. Burns designed and led the study. All authors approved the final manuscript.

