## Supplementary material for "Metagenomic insights into ecosystem function in the microbial mats of Blue Holes, Shark Bay": https://drive.google.com/drive/folders/1hdAbW-tnwMIu5FRgziQOsf89n0OJ2sHQ?usp=sharing

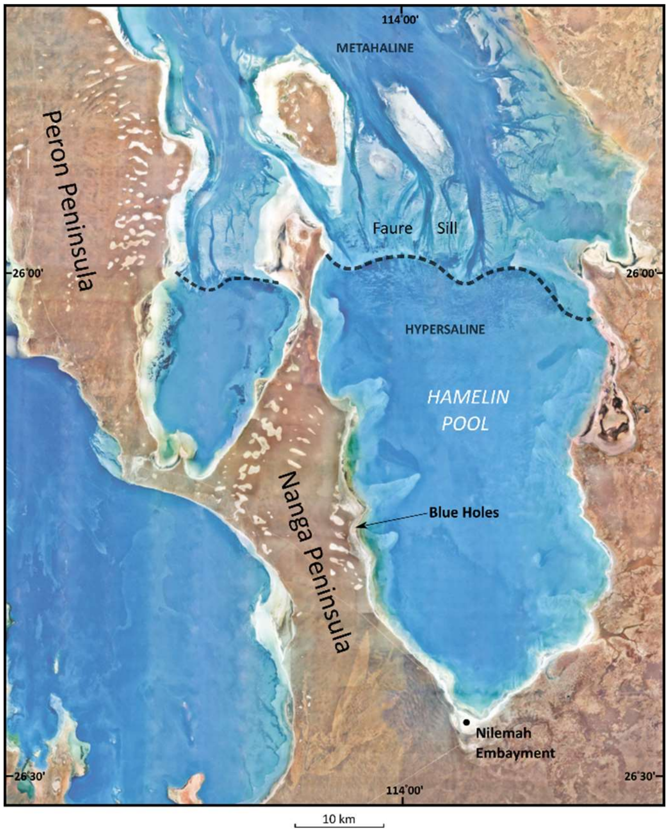


**Fig. S1.**

Map of Hamelin Pool and surrounding areas in Shark Bay. Regions of Shark Bay are metahaline (40–55 PSU) and hypersaline (55-70 PSU). Segmented line displays the low water depth and seagrass sill responsible for restricting the outflow of Hamelin Pool water. Photo provided by the Geological Survey of Western Australia (GSWA).

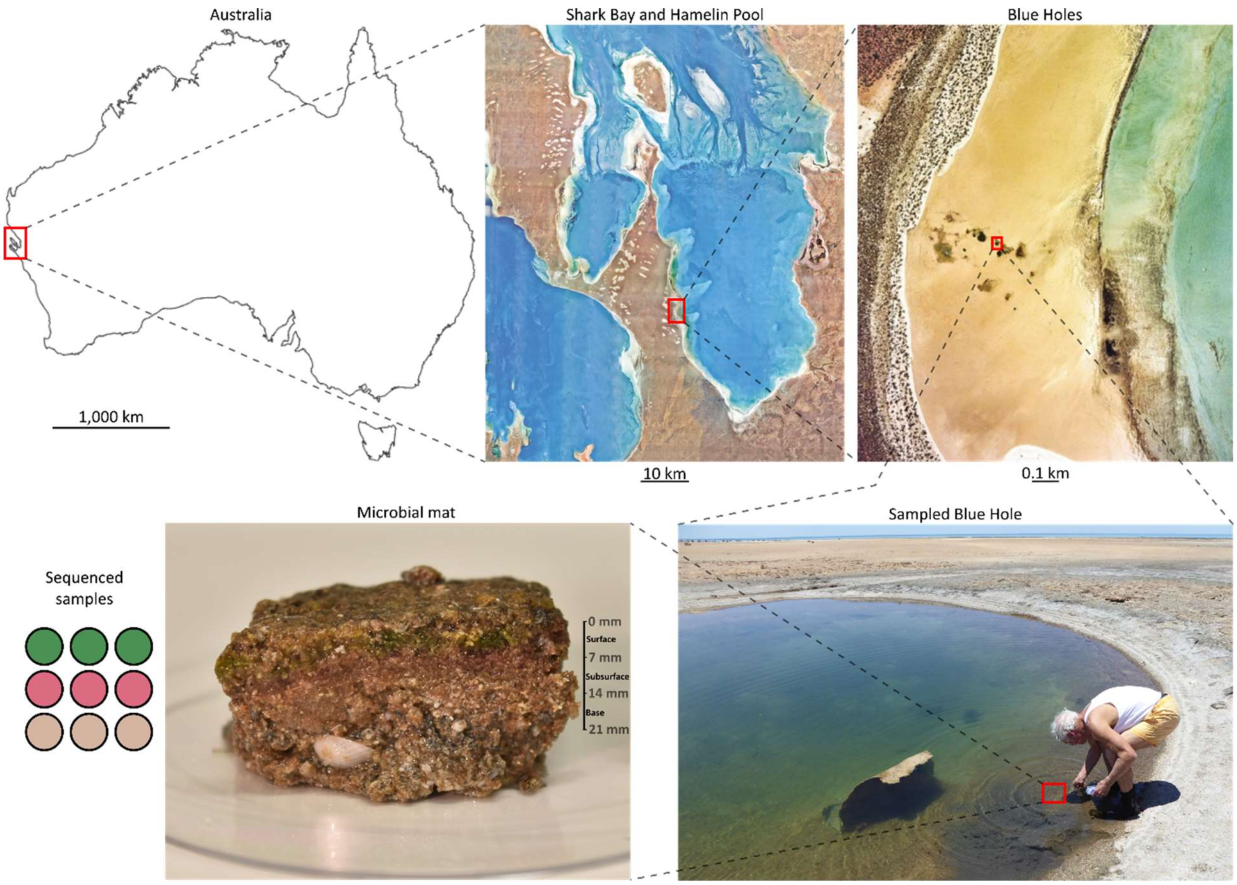


**Fig. S2.**

Geographical and biological perspective of the Blue Holes ecosystem. Aerial photos are adapted from the GSWA. Abnormal structure inconsistent with the hole floor pictured in the bottom right photo is likely the result of a storm separating the surface layer of microbial mat from the underlying sediment. The nine colourised circles represent the nine extractions conducted, three from each layer (surface, subsurface, and base) and reflective of the pigmentation of the mat layer.

A 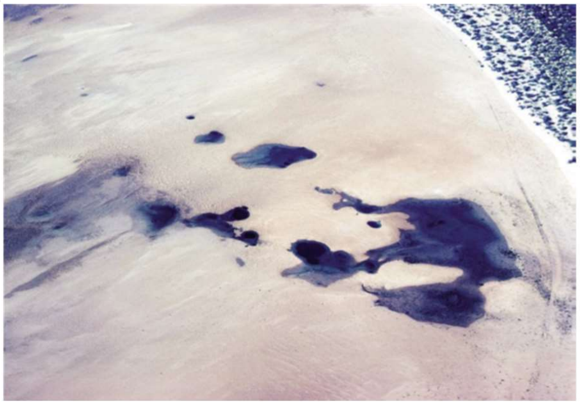 B 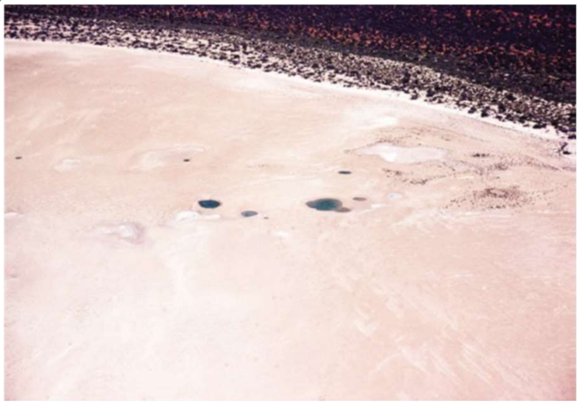

**Fig. S3.**

Aerial photos of Blue Holes showing varying levels of water in the ecosystem. (A) Blue Holes after a period of inundation by either rain or tides, representing high water levels. (B) Blue Holes during a period of high evaporation without refilling by tides or rain, displaying low water levels. Photos provided by the GSWA.

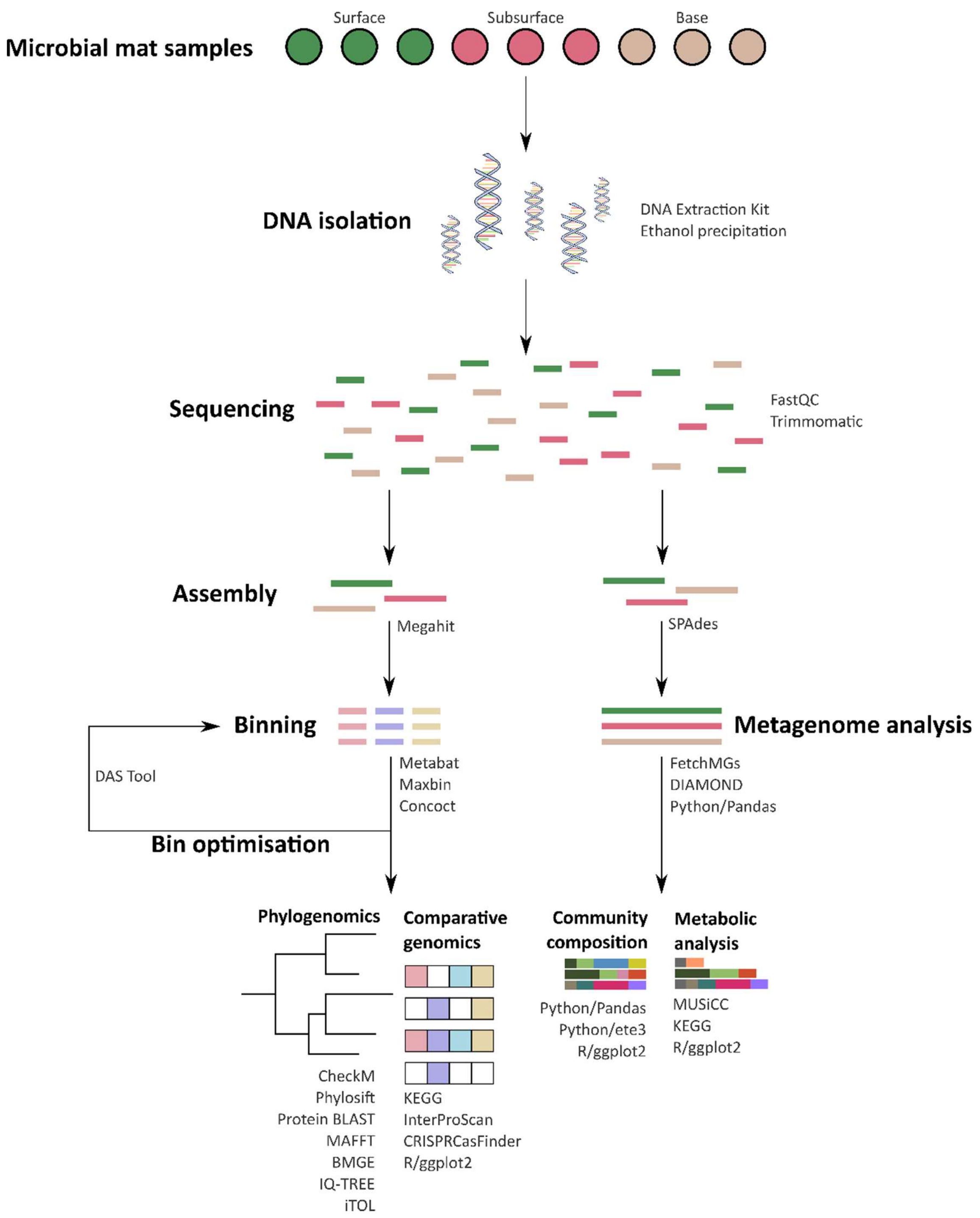


**Fig. S4.**

Workflow of the metagenomic process. Indicated are the step-by-step process with corresponding programs used for analysis. After sequencing, two separate approaches were used to analyse metagenomic data – binning or the creation of metagenome-assembled genomes (MAGs) and metagenome analysis or analysis of the layers of microbial mat.

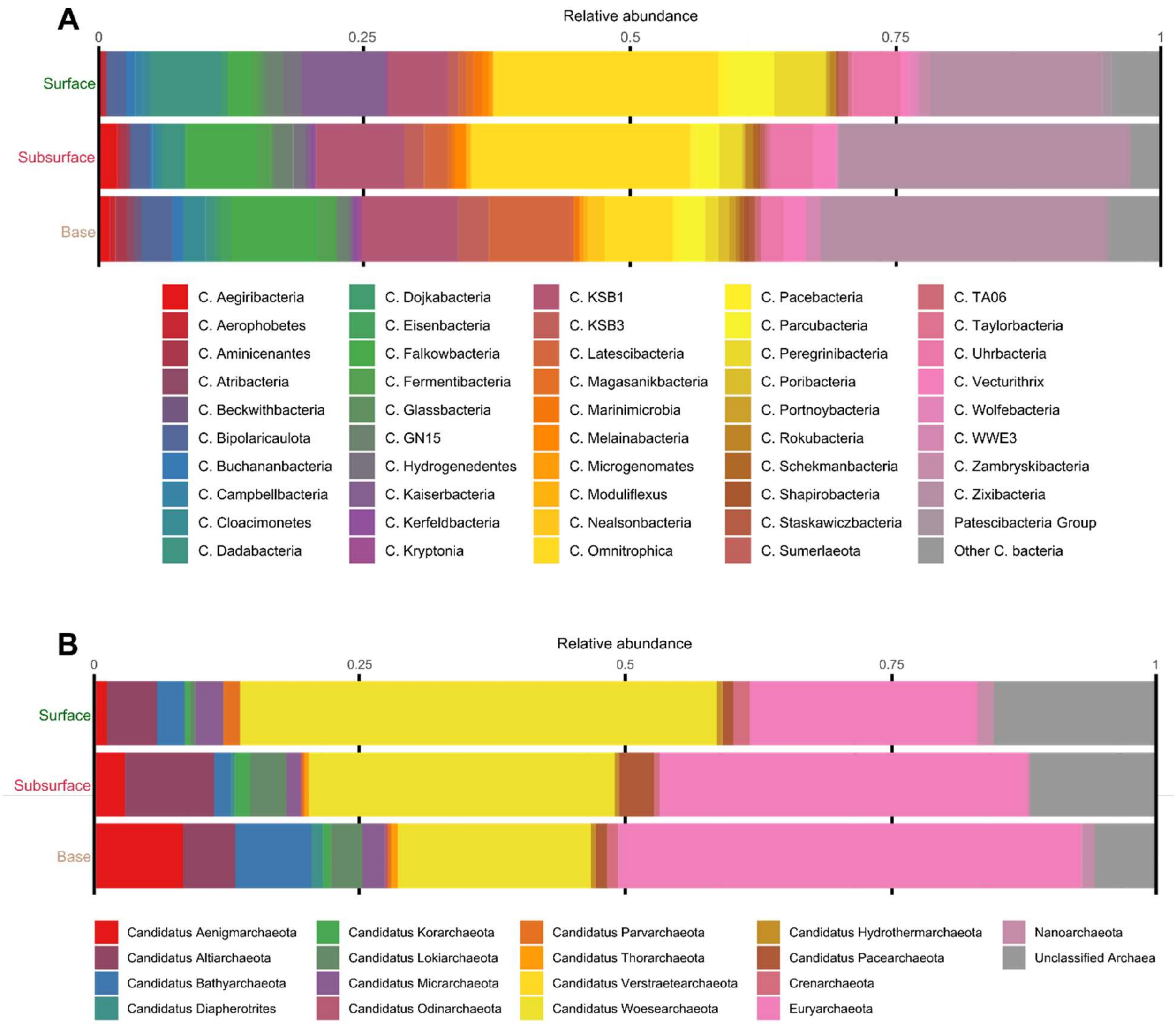


**Fig. S5.**

Relative abundance of bacterial and archaeal phyla found in the Blue Holes microbial mat. A set of 40 universal single-copy genes were used to decipher taxonomy from the metagenomes. (A) Histogram displaying candidate phyla bacteria. (B) Histogram displaying archaeal phyla.

**Supplementary Table 1 (CheckM)**

CheckM, Phylosift, BLASTp, and RNAmmer statistics of Blue Hole MAGs.

**Supplementary Table 2 (BOM Data)**

Shark Bay weather conditions preceding collection. Data acquired from Bureau of Meteorology stations in Shark Bay (Hamelin Station and Hamelin Airport).

**Supplementary Table 3 (Water Chemistry)**

Water chemistry of the water overlying the microbial mats of Blue Holes and Nilemah. Detection limit is the lowest concentration at which an analyte can be detected in a sample with 99% certainty.

**Supplementary Table 4 (Metagenome Stats)**

Assembly and annotation statistics of the three metagenomes derived from the Blue Holes microbial mat layers.

**Supplementary Table 5 (COG Distribution)**

Distribution of universal single copy marker genes (MGs) found in the layers of the microbial mat in Blue Holes. Marker genes were adapted from (Creevey, 2011).

**Supplementary Table 6 (Diversity Relative Abundances)**

Relative abundance values used to create stacked bar charts.

**Supplementary Table 7 (KEGG Genes)**

KEGG genes used as diagnostic of metabolic pathways and cycles.

**Supplementary Table 8 (MUSiCC Relative Abundances)**

Relative abundances combined with MUSiCC-corrected gene counts. RA, relative abundance; MUSiCC, Metagenomic Universal Single-Copy Correction; RA-MUSiCC, relative abundance proportional to MUSiCC value.

**Supplementary Table 9 (MAG Genes Key Functional)**

Genes encoding for key functional nutrient cycles and environmental adaptions. Genes involved in sulphur, nitrogen, carbon, and one-carbon cycles, photosynthesis, phosphate intake, transport, osmotic stress protection, heavy metal and UV resistance, and defences systems are reported. Cells containing “yes” indicates the presence of genes, while “no” shows absence. Genes involved in DISARM have their respective contig locations recorded.
